# Emergence of a Hypervirulent Carbapenem-Resistant *Klebsiella pneumoniae* Co-harbouring a *bla*_NDM-1_-carrying Virulent Plasmid and a *bla*_KPC-2_-carrying Plasmid in an Egyptian Hospital

**DOI:** 10.1101/2021.02.26.433140

**Authors:** Mohamed Abd El-Gawad El-Sayed Ahmed, Yanxian Yang, Yongqiang Yang, Bin Yan, Guanping Chen, Reem Mostafa Hassan, Lan-Lan Zhong, Yuan Chen, Adam P. Roberts, Yiping Wu, Ruowen He, Xiaoxue Liang, Mingyang Qin, Min Dai, Liyan Zhang, Hongyu Li, Yang Fan, Lingqing Xu, Guo-Bao Tian

**Affiliations:** Department of microbiology, School of Basic Medical Science, Xinxiang Medical University, Xinxiang 453003, China; Department of Microbiology, Zhongshan School of Medicine, Sun Yat-sen University, Guangzhou 510080, China; Key Laboratory of Tropical Diseases Control (Sun Yat-sen University), Ministry of Education, Guangzhou 510080, China; Department of Microbiology and Immunology, Faculty of Pharmaceutical Sciences and Drug Manufacturing, Misr University for Science and Technology, Cairo, 6^th^ of October City, Egypt; School of Pharmaceutical Sciences (Shenzhen), Sun Yat-sen University, Guangzhou 510006, China; Department of Neonatal Surgery, Guangzhou Women and Children’s Medical Center, Guangzhou, China; Sun Yat-sen University School of Medicine, Guangzhou 510006, China; Department of Clinical and Chemical Pathology, Faculty of Medicine, Cairo University, Egypt; Department of Tropical Disease Biology, Liverpool School of Tropical Medicine, Pembroke Place, Liverpool, L3 5QA, UK; Centre for Drugs and Diagnostics, Liverpool School of Tropical Medicine, Pembroke Place, Liverpool, L3 5QA, UK; School of Laboratory Medicine, Chengdu Medical College, Chengdu 610500, China; Department of Clinical Laboratory, Guangdong Provincial People’s Hospital / Guangdong Academy of Medical Sciences, Guangzhou, Guangdong, 510080, China; Department of Clinical Laboratory, Sun Yat-Sen Memorial Hospital, Sun Yat-Sen University, Guangzhou 510080, China; Department of Clinical Laboratory, The Sixth Affiliated Hospital of Guangzhou Medical University, Qingyuan People’s Hospital, Qingyuan 511518, China; School of Medicine, Xizang Minzu University, Xianyang, Shaanxi 712082, China

**Keywords:** *Klebsiella pneumoniae*, NDM-1, KPC-2, Hybrid plasmid, Virulent plasmid, Egypt

## Abstract

The emergence of carbapenem-resistant *Klebsiella pneumoniae* (CRKP) isolates in Egyptian hospitals has been reported. However, the genetic basis and the analysis of the plasmids associated with CR-hypervirulent-KP (CR-HvKP) in Egypt are not presented. Therefore, we attempt to decipher the plasmids sequences, which are responsible for transferring the determinants of carbapenem-resistance, particularly the *bla*_NDM-1_ and *bla*_KPC-2_. Out of 34 *K. pneumoniae* isolates collected from two tertiary hospitals in Egypt, 31 were CRKP. Whole-genome sequencing revealed that our isolates were related to 13 different sequence types (STs). The most prevalent ST was ST101, followed by ST383, and ST11. Among the CRKP isolates, one isolate named EBSI036 has been reassessed using Nanopore sequencing. Genetic environment analysis showed that EBSI036 carried 20 antibiotic resistance genes and was identified as CR-HvKP strain, it harboured four plasmids, namely; pEBSI036-1-NDM-VIR, pEBSI036-2-KPC, pEBSI036-3, and pEBSI036-4. The two carbapenemase genes, *bla*_NDM-1_ and *bla*_KPC-2_, were located on plasmids pEBSI036-1-NDM-VIR and pEBSI036-2-KPC, respectively. The IncFIB:IncHI1B hybrid plasmid pEBSI036-1-NDM-VIR also carried some virulence factors, including regulator of the mucoid phenotype (*rmpA*), the regulator of mucoid phenotype 2 (*rmpA2*), and aerobactin (*iucABCD, iutA*). Thus, we set out this study to analyse in-depth the genetic basis of pEBSI036-1-NDM-VIR and pEBSI036-2-KPC plasmids. We reported for the first time a high-risk clone ST11 KL47 serotype of CR-HvKP strain isolated from the blood of a 60-year-old hospitalised female patient from the ICU in a tertiary-care hospital in Egypt, which showed the cohabitation of a novel hybrid plasmid coharbouring the *bla*_NDM-1_ and virulence genes, besides a *bla*_KPC-2_-carrying plasmid.

**IMPORTANCE:** CRKP had been registered in the critical priority tier by the World Health Organization and became a significant menace to public health. Therefore, we set out this study to analyse in-depth the genetic basis of pEBSI036-1-NDM-VIR and pEBSI036-2-KPC plasmids. Herein, we reported for the first time (to the best of our knowledge) a high-risk clone ST11 KL47 serotype of CR-HvKP strain isolated from the blood of a 60-year-old hospitalised female patient in a tertiary-care hospital from the ICU in Egypt, which showed the cohabitation of a novel hybrid plasmid co-harbouring the *bla*_NDM-1_ and virulence genes, besides a *bla*_KPC-2_-carrying plasmid. Herein, the high rate of CRKP might be due to the continuous usage of carbapenems as empirical therapy, besides the failure to implement an antibiotic stewardship program in Egyptian hospitals. Thus, this study serves to alert the contagious disease clinicians to the presence of hypervirulence in CRKP isolates in Egyptian hospitals.

---

Several studies have reported the emergence of carbapenem-resistant *K. pneumoniae* (CRKP) isolates in Egyptian hospitals (1–4); however, to the best of our knowledge, the genetic basis and the analysis of the plasmids associated with CR-hypervirulent-KP (CR-HvKP) in Egypt are not presented. Therefore, we sought to analyse in-depth the genetic basis of pEBSI036-1-NDM-VIR (a novel hybrid plasmid harbouring *bla*_NDM-1_ and virulence genes) and pEBSI036-2-KPC plasmids (a *bla*_KPC-2_-carrying plasmid) which identified from a clinical *K. pneumoniae* strain in Egypt.

A total of 34 nonduplicate *K. pneumoniae* isolates were recovered from the blood of hospitalised patients in two tertiary care hospitals, namely; El-Demerdash hospital (Cairo, Egypt) and National Cancer Institute (Cairo, Egypt) in the period between June 2017 and March 2018 as a part of a study for the monitoring of antimicrobial resistance. Our isolates were selected based on their clinical characteristics, where all of them were primarily identified by VITEK^®^2 and MALDI-TOF MS as *K. pneumoniae* causing bloodstream infections (BSIs), among which 31 were confirmed phenotypically and genotypically as CRKP isolates. Overall, the CRKP isolates were isolated from the blood of 55.9% (19/34) female and 44.1% (15/34) male hospitalised patients, aged from 9 days to 75yr. Minimum inhibitory concentrations of all the 34 isolates were determined for 17 antibiotics using the agar microdilution method according to (CLSI) (5), excepted for tigecycline and colistin using the broth microdilution method according to EUCAST (6). Out of 34 isolates, 91.2% (31/34) were resistant to ertapenem, whereas, 73.5% (25/34) and 61.8% (21/34) were resistant to imipenem and meropenem, respectively. However, all isolates were susceptible to colistin.

All the isolates were assessed by whole-genome sequencing (WGS) using an Illumina HiSeq 2000 platform. *In silico* Multi-locus sequence typing showed that our isolates belong to 13 different Sequence Types (STs). The most prevalent ST was ST101 (13/34, 38.2%), followed by ST383 (5/34, 14.7%). One isolate EBSI036 belongs to ST11, where, ST11 is the dominant ST clone responsible for the prevalence of CRKP worldwide and is considered as an emerging high-risk clone (1, 7–9). According to the clinical data, the *K. pneumoniae* strain EBSI036 was isolated from the blood of a 60-year-old female patient two days after admission to the gastroenterology department of El-Demerdash hospital with symptoms of pneumonia, diarrhoea, and fever. The patient’s symptoms improved following the administration of intravenous ceftriaxone and colistin, and she was discharged from the hospital eight days post-hospitalization. Of note, this strain co-harbours two carbapenemase genes, *bla*_NDM-1_ and *bla*_KPC-2_. Besides, *bla*_SHV-11_, *oqxB, oqxA*, and *fosA6*, were identified in EBSI036 chromosome. The plasmid-associated virulence determinants *rmpA/rmpA2, iucABCD*, and *iutA* in EBSI036 were predicted using the Virulence Factor Database (VFDB; http://www.mgc.ac.cn/VFs/main.htm). EBSI036 was determined as KL47 capsular serotype by using Kaptive software (https://github.com/katholt/Kaptive). The serotype K47 was the most reported type among CRKP infections in Asia (10–12). The virulence level of EBSI036 was confirmed using the *Galleria mellonella* larvae model as previously described (Figure S1) (10, 13). These results revealed that EBSI036 is a CR-hvKP strain.

As the EBSI036 co-harbours two carbapenem genes besides the plasmid-mediated virulence genes, we have further analyzed the characteristics of the related fully sequenced plasmids using a long-read MinION sequencer (Oxford Nanopore Technologies, Oxford, UK). Genomic analysis showed that EBSI036 included a 5,513,124 bp chromosome and four plasmids, namely; pEBSI036-1-NDM-VIR (347 365 bp), pEBSI036-2-KPC (129 869bp), pEBSI036-3 (10 060bp), and pEBSI036-4 (5 596bp) (Table S1). Twenty antimicrobial resistance genes, including six β-lactamases-encoding genes, were identified in EBSI036 using ABRicate version 0.5 (https://github.com/tseemann/abricate) by aligning genome sequences to the ResFinder database. The two carbapenemase genes, bla*NDM-1* and *bla*_KPC-2_, were located on plasmids pEBSI036-1-NDM-VIR and pEBSI036-2-KPC, respectively.

Hybrid plasmids that harbour resistance and virulence genes in a single genetic environment have been reported recently in various *K. pneumoniae* isolates, including the high-risk clones ST23 and ST11 (14–16). Herein, the largest pEBSI036-1-NDM-VIR plasmid belongs to IncFIB: IncHI1B hybrid plasmid. BLASTn showed that pEBSI036-1-NDM-VIR shared >99% identity with plasmid pKpvST383L (CP034201.2), pKpvST147B_virulence (CP040726.1), and p51015_NDM_1 (CP050380.1) with query coverages of 97%-99% (Figure S2). The backbone region of pEBSI036-1-NDM-VIR almost covered the complete sequence of the MDR plasmid pKpvST101_5 with a length of 210 661 bp (CP031372.2) (Figure 1). Most of the remaining sequences (~130 kb) of pEBSI036-1-NDM-VIR were similar to the virulence plasmid pJX6-1 with a length of 228,974 bp (CP064230.1) (Figure 1).

**FIG 1.**
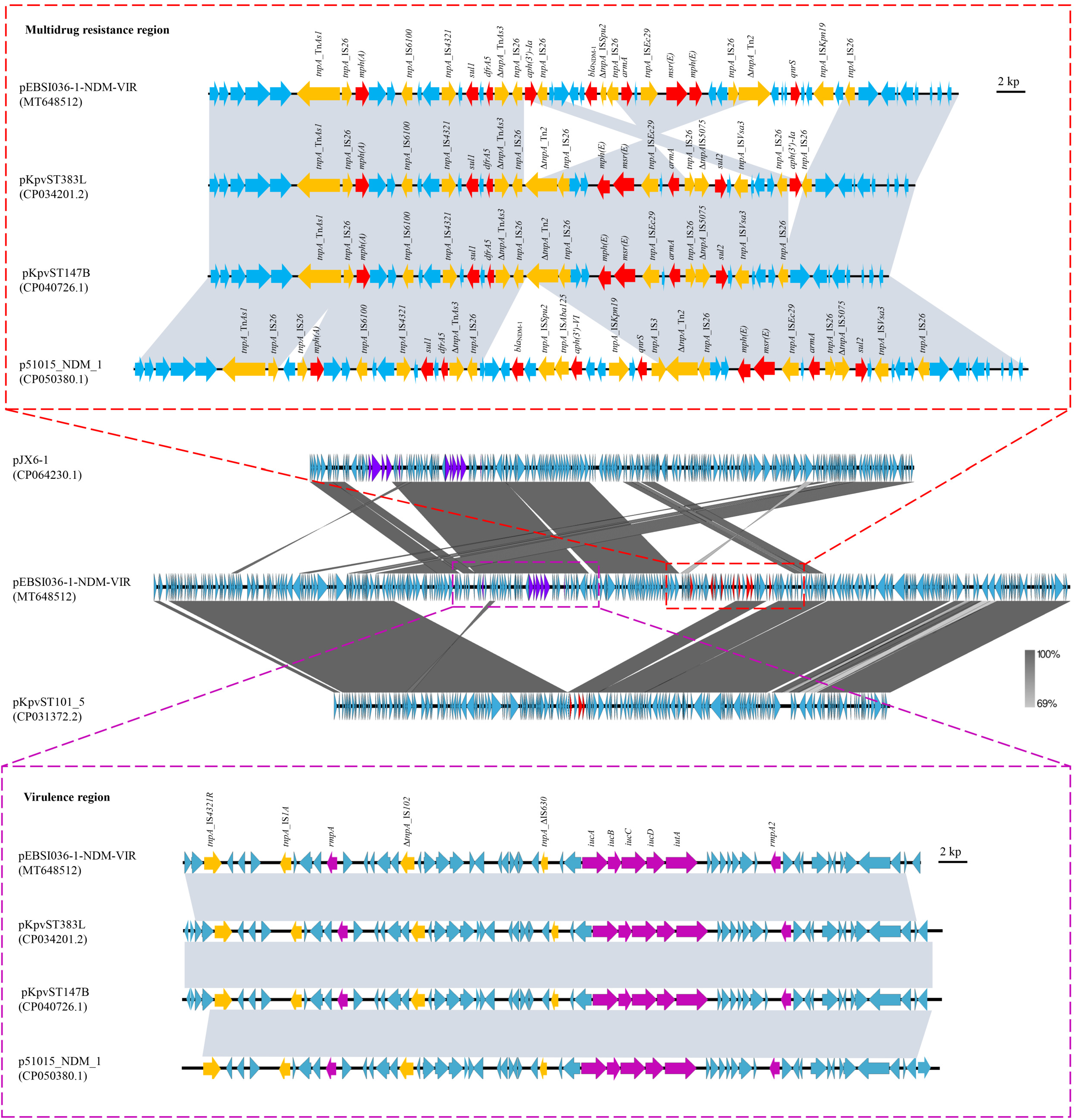
Structure analysis of pEBSI036-1-NDM-VIR. Major structural features of plasmid pEBSI036-1-NDM-VIR were compared with plasmids pKpvST101_5 (GenBank accession number CP031372.2) and pJX6-1 (GenBank accession number CP064230.1). The comparative schematic diagram of resistance region and virulence region in plasmids pEBSI036-1-NDM-VIR, pKpvST383L (GenBank accession number CP034201.2), pKpvST147B_virulence (GenBank accession number CP040726.1) and p51015_NDM_1 (GenBank accession number CP050380.1) were shown respectively. Grey shading indicates shared regions with a high degree of homology. Red and purple represent the antibiotic resistance and virulence genes, respectively, and yellow is the insertion sequences and transposons.

A ~38 kb MDR region in pEBSI036-1-NDM-VIR harboured carbapenemase-encoding gene *bla*_NDM-1_ and another eight resistance genes *mph(A), sul1, dfrA5, aph(3′)-Ia, armA, msr(E), mph(E)*, and *qnrS*. A truncated transposons ΔTn*As1* (Tn*3* family, 6694 bp) and IS*26* elements (IS*6* family, 820 bp) were located upstream of *mph(A)*. The *mph(A)* gene and the downstream complete IS*6100* sequence (Family IS*6*, 880bp) were separated by two ORFs. *sul1* and *dfrA5* were surrounded by IS*4321* (Family IS*110*, 1327bp), ΔTn*As3* (Tn*3* family, 18375 bp), and IS*26* elements. This fragment with 15448 bp containing the above resistance genes was similar to plasmid pKpvST383L (Figure 1). The *aph(3′)-Ia* gene was flanked by IS*26* elements, a similar structure was also found downstream of the resistance region in pKpvST383L. A segment IS*26-armA*-IS*26-msr(E)-mph(E)*-ORF-ORF-IS*26*-ΔTn*2* in pEBSI036-1-NDM-VIR was also found to be identical to the sequence in pKpvST383L with a reversion order. Besides, the *bla*_NDM-1_ and *qnrS* genes were on either side of this fragment, while they were absent in pKpvST383L. By comparing the complete sequences of pEBSI036-1-NDM-VIR and pKpvST383L, it was found that pKpvST383L had another resistance region (26,683 bp) carrying *bla*_NDM-5_ and *bla*_OXA-9_. Compared with plasmid p51015_NDM_1, the resistance region of plasmid pEBSI036-1-NDM-VIR lacked the *aph(3′)-VI* and *sul2* genes (Figure 1). It is worth noting that in pEBSI036-1-NDM-VIR, the multidrug-resistant region contained six IS*26* elements and other transposon elements. Some studies demonstrated that the resistance loci containing IS*26* can be hotspots for the capture of further resistance genes to constitute a novel multi-drug resistant region (17).

A group of virulence genes was detected in pEBSI036-1-NDM-VIR; *rmpA* and *rmpA2*, which are commonly attributed to the hypermucoviscous phenotype of *K. pneumoniae* and the *iucABCD* and *iutA*, associating with virulence, with increased colonisation and infection producing capabilities (18). The ~39 kb region harbouring virulence genes exhibited high similarity (99.9% identity and 98% query coverage) with pKpvST383L, pKpvST147B_virulence, and p51015_NDM_1 (Figure 1).

The pEBSI036-2-KPC plasmid was determined to be an IncR: IncFII-type plasmid. Comparative analysis showed that pEBSI036-2-KPC had 98%-99% query coverages and 99.9% nucleotide identity with the following plasmids, pKP19-2029-KPC2 (CP047161.1), p69-2 (CP025458.1), and p16HN-263_KPC (CP045264.1). The pEBSI036-2-KPC plasmid carried carbapenemase-encoding gene *bla*_KPC-2_ and three β-lactamases-encoding genes; *bla*_CTX-M-65_, *bla*_TEM-1B_, and *bla*_SHV-12_. The pEBSI036-2-KPC plasmid had additional resistance genes; *catA2, fosA3*, and *rmtB*. These resistance genes were located in two main resistance regions (Figure 2). The *bla*_KPC-2_ and *bla*_SHV-12_ genes were separated by sequence ΔTn*As1*-IS*26*-ΔTn*3*-IS*Kpn27*, and IS*Kpn6* was located downstream of *bla*_KPC-2_. The downstream of *bla*_KPC-2_ contained a *mer* operon responsible for mercuric resistance and transposons elements (ΔTn*As1*-IS*26*-ΔTn*As3*). This segment carrying *bla*_KPC-2_, *bla*_SHV-12_, and a *mer* operon, was highly similar to other plasmids such as pKP1034 (19). There was another MDR region (15 254 bp) that consisted of *bla*_CTX-M-65_, *fosA3, bla*_TEM-1B_, and *rmtB* genes, and five IS*26* fragments (Figure 2). The basic structure of pEBSI036-2-KPC is similar to plasmid pKPC2_040035 (CP028796.1) (99.98% identity and 88% query coverage), except for two regions. One 10,379-10,795 locus carried *fosA3*, which was flanked by IS*26*. The other region contained ΔIS*Cfr3*-IS*Kpn26*-IS*26-catA2*-IS*26*-IS*5075*-ΔTn*3*-IS*26* structure with a length of 15,042bp, which was the same as plasmid p3_L382 (CP033962.1) with 100% query coverage and 99.99% nucleotide identity (Figure 2). Both *fosA3* and *catA2* were flanked by IS*26* as previously reported (19, 20). That evidence emphasizes the role of insertion elements such as IS*26* in insertion and deletion of resistance genes again.

**FIG 2.**
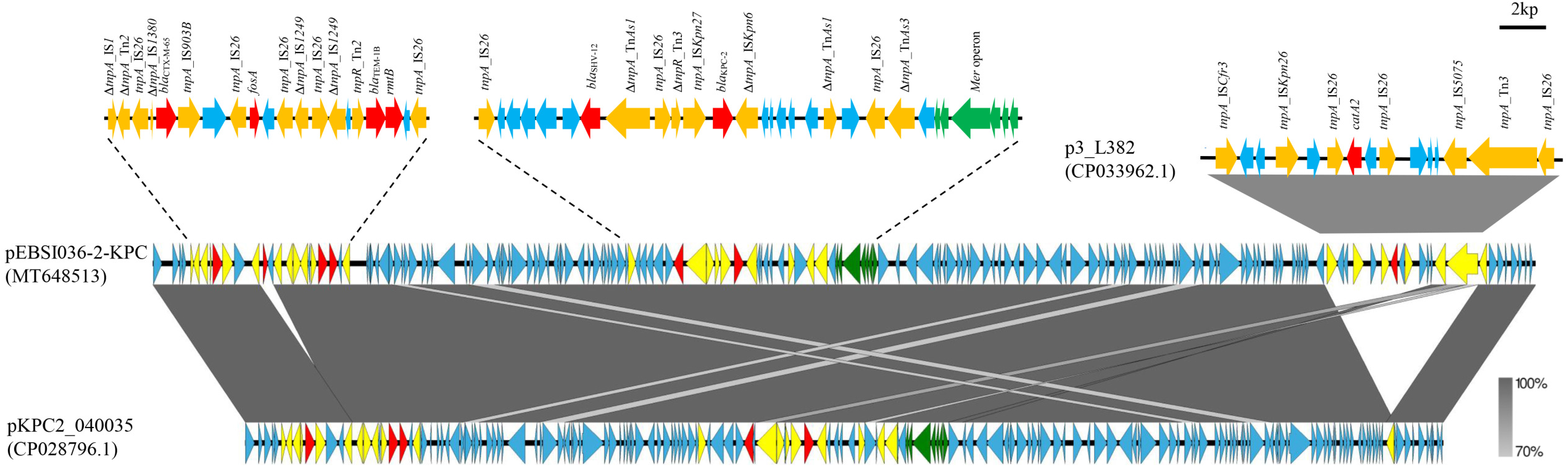
Sequence alignment analysis among plasmids pEBSI036-2-KPC, pKPC2_040035 (GenBank accession number CP028796.1) and p3_L382 (GenBank accession number CP033962.1). Red and green represent the antibiotic resistance and heavy metal resistance genes, respectively. And yellow is the insertion sequences and transposons.

In conclusion, to the best of our knowledge, this is the first report of a high-risk clone ST11-KL47 of CR-HvKP strain isolated from the blood of a patient from the ICU in Egypt, co-harbouring two plasmids, one is a novel hybrid plasmid harbouring the carbapenemase gene *bla*_NDM-1_ and virulence genes, and the other is a *bla*_KPC-2_-carrying plasmid. Further countrywide surveillance studies are needed to elucidate the rate of prevalence of this high-risk clone in Egypt and its burden on hospital-acquired infections.

## Accession numbers

The sequences of the plasmids pEBSI036-1-NDM-VIR and pEBSI036-2-KPC were deposited in GenBank with accession numbers MT648512 and MT648513.

## ACKNOWLEDGMENTS

This work was supported by the National Natural Science Foundation of China (grant numbers 82061128001, 81722030, 81830103, 81902123), National Key Research and Development Program (grant number 2017ZX10302301), Guangdong Natural Science Foundation (grant number 2017A030306012), Project of high-level health teams of Zhuhai at 2018 (The Innovation Team for Antimicrobial Resistance and Clinical Infection), 111 Project (grant number B12003), Open project of Key Laboratory of Tropical Disease Control (Sun Yat-sen University), Ministry of Education (grant numbers 2020kfkt04,2020kfkt07) and China Postdoctoral Science Foundation (grant number 2019M653192), Science, Technology & Innovation Commission of Shenzhen Municipality (grant numberJCYJ20190807151601699), the Science and Technology Planning Project of Guangdong (grant number2017A020215017), Qingyuan People’s Hospital Medical Scientific Research Fund Project (grant number 20190209), Guangdong Provincial Bureau of Traditional Chinese Medicine research fund (grant number 20201407).

## Supplementary Materials

**Table S1** Overall features of the *K. pneumoniae* EBSI036 genome

**Table S2** Putative virulence genes detected on the *K. pneumoniae* EBSI036 chromosome

**FIG S1** Virulence potential of *K. pneumoniae* strain EBSI036 as depicted in a *Galleria mellonella* infection model with an inoculum of 1 × 10^4^CFU.

**FIG S2** Sequence alignment analysis among plasmids pEBSI036-1-NDM-VIR and pKpvST383L (GenBank accession number CP034201.2), pKpvST147B_virulence (GenBank accession number CP040726.1) and p51015_NDM_1 (GenBank accession number CP050380.1).

